# Mitochondrial transplantation and its impact on infectious disease progression: a pilot study

**DOI:** 10.1101/2023.09.08.556161

**Authors:** Tom Benson, Samir P. Patel, Benedict C. Albensi, Vinit B. Mahajan, Aida Adlimoghaddam, Sergey Sikora, Hiroshi Saito

## Abstract

**BACKGROUND:** Mitochondrial transplantation has recently gained
prominence as a novel technique primarily focused on addressing ischemiareperfusion injuries and rare mitochondrial mutation diseases. Platelets, abundant in the bloodstream, play a crucial role in immune function. Upon activation, platelets release mitochondria encapsulated within extracellular vesicles, here referred to as “mitlets”. These mitlets exhibit a preference for being internalized by immune cells circulating in the bloodstream, enhancing their cellular energetics. Herein, we hypothesized that the transplantation of mitlets between young animals and aged animals may exert a significant influence on the progression of infectious diseases.

**STUDY DESIGN AND METHODS:** In this study, murine models of Influenza H1N1 infection and sepsis were employed to investigate disease dynamics. Specifically, mitlets isolated from young and healthy mice were transplanted into cohorts of mice of the same age afflicted by H1N1 infection, or into aged mice subjected to polymicrobial infection and sepsis. Survival outcomes and the quantification of cytokine levels were assessed across experimental groups to elucidate the potential therapeutic effects of mitlet transplantation.

**RESULTS:** In the matched-age H1N1 infection model, as predicted, mitlet transplantation did not yield a statistically significant improvement in survival, although it did show a trend towards a reduction in the circulating inflammatory cytokine burden. In the young-to-old sepsis model, the transplantation of mitlets was associated with a significant enhancement in survival rates and a substantial reduction in bacterial loads and circulating cytokine levels.

**DISCUSSION:** Our findings suggest that mitochondrial transplantation may constitute a safe and promising avenue for enhancing the immune system’s capacity to counter infectious threats. This pilot investigation sets the stage for further exploration. It is plausible that in the future, immune senescence resulting from diminished mitochondrial energy production
could be ameliorated through such transplantation interventions. As a consequence, this approach holds substantial potential as a novel immunotherapeutic strategy for the management of infectious diseases.

## 1 INTRODUCTION

Mitochondria produce the majority of cellular energy and, therefore, the energy for immune system function. Mitochondrial energy production declines with age, and this decline is more pronounced in primates and humans compared to mice. For example, the energy decline in retinal mitochondria of 14-year-old)\ primates is 70 percent,, while the same decline measured in 12-month-old mice is 30 percent.^85^ Recent work has found that mitochondria are frequently transferred between cells via stem cells and extracellular vesicles^21^, and researchers have tested many techniques for effecting such transfers from an exogenous source – isolating and transplanting the mitochondria from cell to cell or from animal to animal.^72,87,88^ Platelets are the most common form of immune system blood cell.^27^ Mitochondria can be found in great numbers in the bloodstream, emitted inside extracellular vesicles by platelets during activation or at end-of-life.^14^ Research has found that such platelet-derived mitochondrial vesicles (“mitlets”) are quickly and preferentially absorbed by other immune cells in the bloodstream, including white blood cells and other platelets.^11^

Mitochondrial transplantation is an emerging technique in which exogenous mitochondria, isolated from either the patient or a donor or grown in external cell cultures, are transplanted into organs to restore cellular energetics and cure injuries. This technique until now has been tested primarily on ischemia-reperfusion injuries, organ transplants, and rare mitochondrial mutation diseases but has not been explored for adult age-related or chronic diseases.

Historically, the elderly are more prone to death and disability from infection. The age-related mechanisms that reduce the effectiveness of the immune system are not fully understood. We hypothesized that the decline of the human immune system, and thus the increased levels of mortality and morbidity in the elderly due to infectious disease, is primarily due to reduced energy production by mitochondria inside white blood cells and immune cells. The functions of white blood cells are energy-intensive, particularly during infection. If they suffer reduced energy production due to age, then it follows that they will be unable to perform their functions effectively during a serious infection. Aged white blood cells move slower, act slower, and display a reduced ability to digest or kill pathogens compared to younger white blood cells.^1^

We tested this hypothesis by transplanting mitlets containing mitochondria. We isolated the mitlets from the blood of young animals and then transplanted them into cohorts of young and aged animals with induced viral or bacterial infections. We performed similar transplants for mitochondria that were isolated from mouse liver, not encased in vesicles. We then ascertained survival, blood pathogen load, and cytokine levels for these mitochondrial types and a variety of delivery mechanisms.

## 2 METHODS

### 2.1 Facilities involved

Experiments below were accomplished at several locations. Charles River Labs in Bristol, England performed experiments using mice with an H1N1 influenza virus. The University of Kentucky, USA performed experiments using mice with an induced sepsis infection. Additional mitlet preparation and testing was performed by AA at St. Boniface Hospital, MB, Canada.

### 2.2 Ethics

All experimental procedures at the University of Kentucky were approved by the Institutional Animal Care and Use Committee. Animal handling techniques in these studies were performed as described and approved in our Animal Use Protocol #2009-0541 and were in accordance with the National Institutes of Health guidelines for ethical treatment. All experimental procedures at Charles River Labs in Bristol, England, were approved by the Institutional Animal Care and Use Committee. Animal handling techniques in these studies were performed as described following commonly-accepted guidelines for ethical treatment.

### 2.3 Mice for sepsis experiments, University of Kentucky

For these experiments, middle-aged (13-month-old) female C57BL/6 mice with jugular vein catheterization were obtained from Charles River Laboratories, Watsonville, CA. A Vascular Access Button (VAB, Intech Laboratories, Inc) was attached to the subcutaneous end of the catheter, allowing exogenous fluids infusion repeatedly. Animals were maintained in the Division of Laboratory Animal Resources at the University of Kentucky in pressurized intra-ventilated cages housed five animals per cage under controlled temperature (21 – 23°C), humidity (30% – 70%), and lighting (14/10 light/dark cycle) with free access to drinking water and chow (Teklad Global No. 2918, 18% Protein Rodent Diet, Madison, Wis). All animals were acclimated for at least 7 days prior to experimentation to eliminate the influence of transportation stress.

### 2.4 Induction of sepsis in mice at University of Kentucky

Experimental polymicrobial abdominal sepsis was induced by bolus injection of cecal slurry (CS) as we previously described in Steel et al.^66.^ Survival and the health of each mouse was monitored multiple times daily for 14 days. A small amount of blood samples was collected aseptically from the tail vein of each mouse 12 hours after sepsis induction. Each blood sample (10-μL) was quickly diluted with 9-folds volume (90-μL) of 1x citrate solution (0.32% sodium citrate in 0.9% sodium chloride) and plated onto an agar plate as we described (Steel et al ^66^). Anaerobic bacteria colonies were counted after incubating these agar plates for 2 days using the GasPak anaerobic gas generating pouch system (BD Biosciences).

### 2.6 Mice for influenza virus experiments, Charles River Labs, Bristol, England

Adult female BALB/c mice from internal stock were randomly allocated and allowed to acclimatize for one week.

### 2.7 Induction of H1N1 virus in mice at Charles River Labs, Bristol, England

On Day 0, animals were challenged with A/Puerto/Rico/8 (H1N1)(PR8) and monitored daily for clinical signs of ill-health until termination on Day 7. In-life bleeds were taken on Day -1 (pre-treatment) and Day 3 (post-treatment) and processed to plasma for Luminex analysis of cytokine levels. At termination, bronchoalveolar lavage (BAL) and blood was taken. Blood was processed to plasma and serum. BAL fluid and plasma samples were analyzed for cytokine levels using Luminex and serum was analyzed in a HAI assay to indicate the levels of functional anti-viral antibodies present.

### 2.8 Platelet isolation and preparation of platelet extracellular vesicles (mitlets) at University of Kentucky for sepsis experiments

We used the same approach described by Marcoux et al. to isolate platelets and mitlets^32^. In brief, C57BL/6 2-month-old mouse blood preserved with ACD 20% on ice was purchased from BioIVT Inc., Westbury, NY. Platelets were isolated from blood, and then washed with Tyrode’s Buffer pH 7.4 and resuspended at 1x10^8^ cells/mL in Tyrode’s Buffer pH 7.4 with 5 mM CaCl_2_. Platelets were stimulated overnight (16h) at room temperature with thrombin. 10 mM EDTA was added to stop the stimulation, then stimulated platelets were centrifuged at 300 g for 5 min to remove remnant platelets or cells. The resulting mix is concentrated by ultracentrifugation at 18,000 *g* for 60 min at 18°C, which isolates approximately 98% of the mitlets according to quantifications of protein contents^3.2^ The mitlet pellet was resuspended in filtered phosphate-buffered saline (PBS) pH 7.4. The resulting mitlets in PBS were chilled to 4 degrees C for up to 1 week prior to administration.

### 2.9 Platelet isolation and preparation of platelet extracellular vesicles (mitlets) performed at St. Boniface Hospital in support of Bristol H1N1 virus experiments

We used the same approach described by Marcoux et al. to isolate platelets and mitlets^32^. In brief, 2-month-old male or female BALB/C mouse blood preserved with ACD 20% at room temperature was purchased from BioIVT Inc., Westbury, NY. Platelets were isolated from blood, and then washed with Tyrode’s Buffer pH 7.4 and resuspended at 1x10^8^ cells/mL in Tyrode’s Buffer pH 7.4 with 5 mM CaCl_2_. Platelets were stimulated overnight (16h) at room temperature with thrombin. 10 mM EDTA was added to stop the stimulation, then stimulated platelets were centrifuged at 300 g for 5 min to remove remnant platelets or cells. The resulting mix is concentrated by ultracentrifugation at 18,000 *g* for 60 min at 18°C, which isolates approximately 98% of the mitlets according to quantifications of protein contents.^3.2^ The mitlet pellet was resuspended in filtered phosphate-buffered saline (PBS) pH 7.4. 50% of the resulting mitlet/IPBS mixture was mixed with an additional cryopreservative and chilled to 4 degrees C. The other 50% of the mitlets were left in PBS and chilled to 4 degrees C. All chilled mitlets were shipped on ice by medical courier to the Charles River Labs, Bristol, England location.

### 2.10 Liver mitochondrial preparation for sepsis tests at University of Kentucky

Mitochondria were isolated from young (2-month-old) healthy C57BL/6 mouse liver from internal stock, as described before^32^. In summary, naive anesthetized mice under isoflurane were euthanized by cervical dislocation, and livers were removed and placed in ice cold homogenization buffer. Livers were transferred into a Dounce tissue homogenizer along with 5 ml ice cold homogenizing buffer (300 mM sucrose, 10 mM K-HEPES, and 1 mM K-EGTA (pH 7.2)). The tissue was gently homogenized for 60 seconds. 250μL of Subtilisin A (96.61μM) was added to the homogenate and mixed by inversion. This was then incubated on ice for 10 minutes. Using a pre-wet 40μm filter, filter the homogenate into a 50mL falcon on ice. This step was repeated with a 40μm filter and then a 10μm filter. The filtrate was then transferred to 1.5mL Eppendorfs and centrifuged at 9,000 xg for 10 minutes at 4°C. The supernatant was discarded, and the pellet was resuspended in 1mL ice-cold PBS. The concentrated filtrate was then filtered through a 1.2μm and 0.8μm filter. A protein assay was performed to confirm the concentration of the mitochondria. This was adjusted to 10mg/kg and handed to the in vivo team on ice for administration.

### 2.11 Liver mitochondrial preparation for H1N1 Viral Tests at CRL Bristol, England

Mitochondria were isolated from young (2-month-old) healthy C57BL/6 mouse liver from internal stock, as described before^32^. In summary, naive anesthetized mice under isoflurane were euthanized by cervical dislocation, and livers were removed and placed in ice cold homogenization buffer. Livers were transferred into a Dounce tissue homogenizer along with 5 ml ice cold homogenizing buffer (300 mM sucrose, 10 mM K-HEPES, and 1 mM K-EGTA (pH 7.2)). The tissue was gently homogenized for 60 seconds. 250μL of Subtilisin A (96.61μM) was added to the homogenate and mixed by inversion. This was then incubated on ice for 10 minutes. Using a pre-wet 40μm filter, filter the homogenate into a 50mL falcon on ice. This step was repeated with a 40μm filter and then a 10μm filter. The filtrate was then transferred to 1.5mL Eppendorfs and centrifuged at 9,000 xg for 10 minutes at 4°C. The supernatant was discarded, and the pellet was resuspended in 1mL ice-cold PBS. The concentrated filtrate was then filtered through a 1.2μm and 0.8μm filter. A protein assay was performed to confirm the concentration of the mitochondria. This was adjusted to 10mg/kg and handed to the in vivo team on ice for administration.

### 2.12 Statistical analysis and software at University of Kentucky

Survival curves were analyzed by Kaplan–Meier LogRank test. Data for two-group comparisons were analyzed by Student t test. When multiple comparisons were made, the Shapiro–Wilk normally test was run. If the data passed the normality test, one-way ANOVA and Holm–Sidak post-hoc test were used to analyze the data. Alternatively, when the data was not normally distributed, the Kruskal–Wallis test and Dunn post-hoc test were used. In instances where one group was assessed multiple times (i.e., bacteria load), repeated-measures one-way ANOVA was used, and when multiple groups were assessed multiple times (i.e., body temperature data), repeated-measures two-way ANOVA was used and the Holm–Sidak post-hoc test was run. All data are expressed as means and standard deviations and p <0.05 was considered statistically significant.

### 2.13 Statistical analysis and software at Charles River Labs, Bristol, England

Samples were analyzed individually at a 1 in 4 dilution. Each dot represents the group mean + SEM at each time point. The functional Lower Limit of Quantification (LLOQ) (3.34 pg/mL) is plotted as a dotted black line. Values <LLOQ are plotted as 0 pg/mL. Statistical significance analysis was attempted using a Mixed-effects model, however due to values <LLOQ a p value could not be calculated. Normality was determined by a Shapiro-Wilk test. Statistical significance between the Naïve (uninfected) and Vehicle group was determined by a Mann-Whitney T test. Statistical significance between the vehicle, oseltamivir, Mitlets and Liver mitochondria-treated groups was determined using a Kruskal-Wallis test followed by Dunn’s multiple comparison test, using the vehicle group as the comparator. No differences between the groups in any tests were found to be statistically significant.

## 3 RESULTS

### 3.1 Mitlets transplanted into age-matched mice with H1N1 influenza slightly improved survival and cytokines

(Charles River Labs, Bristol, England location). Liver isolated mitochondria had no noticeable impact. Adult female BALB/c mice were randomly allocated and allowed to acclimatize for one week. Each immune-assay readout was performed individually. Readouts were: Bodyweights, Clinical Scores, Survival, Cytokine levels measured using Luminex: 5-plex testing IL-6, IL-10, IL-17, IFN-ƴ and TNFα Haemagglutination inhibition assay (HAI). Compounds were administered by tail-vein injection, according to the table below:

| Groups | N= | Treatments |  |  | Intervention |
| --- | --- | --- | --- | --- | --- |
|  |  | Dose | Route | Regimen |  |
| Naive | 5 | N/A | IV, 120 μL | Day -1, Day 1 and Day 3 (x3) | N/A |
| Vehicle control | 5 | N/A | IV, 120 μL | Day -1, Day 1 and Day 3 (x3) | Day 0: IN admin of PR8 H1N1 influenza virus |
| Oseltamivir | 5 | 20mg/kg | PO | SID from Day -1 to end |  |
| Mitlets (in PBS) | 5 | N/A | IV, 120 μL | Day -1, Day 1 and Day 3 (x3) | |
| Mitlets (in Cryopreservative) | 5 | N/A | IV, 120 μL | Day -1, Day 1 and Day 3 (x3) | |
| Liver Mitochondria | 10 | 10mg/kg | IV, 120 μL | Day -1, Day 1 and Day 3 (x3) | |

As predicted, mitlet injections slightly improved survival and reduced cytokines, but not with statistical significance. Isolated liver mitochondria showed no measurable effects.

### 3.2 Mitlets transplanted from 2-month mice into 13-month mice, induced with sepsis, significantly improved survival and cytokines

(University of Kentucky location). 13-month-old C57BL/6 mice were surgically modified by the vendor to add a VAB (injection port) in the jugular vein for easy injection. Mice were allowed to acclimate in cages for 1 week after receipt, then infected with Sepsis per long-established protocol^66^. Mice were injected according to the following table:

| Groups | N= | Treatments |  |  | Intervention |
| --- | --- | --- | --- | --- | --- |
|  |  | Dose | Route | Regimen |  |
| Vehicle control with mitlets | 8 | N/A | IV, 120 μL | Day -1, Day 1, Day 2 & Day 3 (x4) | Day 0: induction of Sepsis via Cecal Slurry injection |
| Mitlets (in PBS) | 8 | N/A | IV, 120 μL | Day -1, Day 1 and Day 3 (x3) | |
| Vehicle control with liver mitochondria | 3 | 10mg/kg | IV, 120 μL | Day -1, Day 1 and Day 3 (x3) | |
| Liver mitochondria | 5 | 10mg/kg | IV, 120 μL | Day -1, Day 1 and Day 3 (x3) | |

They were injected at -1, 1, 2, and 3 days. The bacterial count and cytokines were collected at -2, 1, and 3 days. The study was duplicated several months later with identical results. An additional 3 treatments on Days 3, 4, and 5 were performed or Exp 2. In the first experiment, 2 mice died during injection, probably because of poor handling, and were removed from the experiment.

Mitlet treatments significantly prevented early death of mice one week after sepsis induction (P<0.01). Mitlet treatments (by Day 2 - 5) delayed but did not prevent later death. Mitlet treatment significantly reduced blood bacteria several folds but didn’t eliminate it completely, and reduced IL-6 cytokine level ∼ 60%. The effect of mitlet treatment was temporary and declined a few days after the last injection. In contrast to mitlets, injections of mitochondria isolated from the liver had no protective effect – mice in these cases died at the same time as negative controls.

## 4 DISCUSSION

In our study, we transplanted mitochondria at two different locations using two separate teams, at Charles River Labs in Bristol, England (Bristol) and the lab of Hiroshi Saito at the University of Kentucky (Kentucky). At Bristol, we transplanted same-age mitochondria for BALB/C mice, 2-3 months old. Recipient mice were dosed with an H1N1 flu virus. We used same-age donor mitochondria of three types: mitochondria isolated directly from the livers of donor mice and mitlets isolated from the blood of BALB/C mice in another lab in Winnipeg, Canada, either preserved in a cryopreservative solution or PBS, chilled to 4° C, and shipped by courier to Bristol. All three of these were injected into recipient mice three times over seven days. We did not expect to see a beneficial effect in the Bristol experiments because the mitochondria were from animals of the same age; however, there was a small noticeable effect, albeit not enough to reach statistical significance. For the Kentucky tests, we injected mitochondrial test products from 2-month-old C57BL/6 mice into aged 13-month-old C57BL/6 mice, which had been given sepsis via cecal slurry induction. The first injections were mitochondria isolated from the liver of the same-aged mice and injected immediately, which showed no results. The second injections were mitlets isolated from the blood of donor animals, chilled in PBS to 4° C, and then injected three different times over the course of the experiment, which showed highly significant improvements (p < 0.05). We duplicated the successful mitlet experiments later with the same results.

These experiments, taken together, correspond accurately to our predictions and to the in-vitro work done in parallel at our partner lab at the University of Laval Quebec. ^68,32^ We expected to see that mitochondria transplanted from younger to older mice would create a much largers beneficial effect. We expected that mitlets would provide a robust response while liver-isolated mitochondria would produce little or no response because mitlets are preferentially targeted to immune cells, while liver mitochondria, with no vesicle and no receptors, become scattered into different tissues.

### The benefits of mitlets are more pronounced in aged mice

Aged mice have mitochondria that are more dysfunctional compared to young mice. This has been shown in prior studies that looked at the energy production level of mitochondria in aged animals versus young animals.^85,86^ According to the “Mitochondrial Theory of Aging^83^,” mitochondria start as highly functional at birth, but their functionality and performance decline steadily over the years, causing an inexorable decline in the ATP energy produced in each cell. As a result, the total energy available to a given organ or sub-system of the body, in this case the immune system, declines with age.^84^ A similar decline can also be seen in children with mitochondrial mutation diseases such as Leigh syndrome or Kearns-Sayre syndrome. They are born with mutated and dysfunctional mitochondria from birth and are subject to a large variety of morbidity and mortality due to insufficient cellular energetics.^63,64^ These children are most affected when the ratio of dysfunctional mitochondria exceeds a certain threshold, generally 60-70%.^65^ Thus, we see that mitochondrial energy production is a measurable and predictable number, which can be seen to have predictable consequences for the human body. In a similar way, mitochondrial energy production declines due to age, in some cases 70% in the elderly primate retinas.^85^ We and other researchers hypothesize that these age-related energy reductions lead to cellular failure, lower activity, less performance, accumulation of waste products, and other effects that result in a large part of biological aging. In this work, we tested the ability of white blood cells to combat infection, which is reduced in the 13-month-old mice used in Kentucky. We injected mitlets from young animals, which are absorbed into the white blood cells and used as supplemental energy sources to support the increased energy demand associated with their antimicrobial functions, such as phagocytosis, degranulation, and NETosis. Thus, we are, in essence, making these white blood cells temporarily younger, or we may instead say, we are making them temporarily *act* younger.

### Lesser results for same-age animals

We weren’t sure if we would see measurable results for same-age animals, because there is no difference in the activity level and the energy production of mitochondria in the donor mitochondria versus the wild-type mitochondria in the white blood cells of the recipient animals. It was interesting that the transfusion of mitlets produced a modest effect. We interpret this to mean that the recipient white blood cells, although they received mitochondria of approximately the same functionality as those they already had, nevertheless benefited by getting additional mitochondria. According to our *in-vivo* studies^68^, white blood cells have a certain “starting complement” of existing mitochondria. In the middle of an infection, the white blood cells might have the capacity to accept additional mitochondria – again, corresponding to our earlier analogy of mitochondria as a type of spare battery pack. We helped the white blood cells not by giving them *higher-quality* mitochondria, but simply by giving them *more* mitochondria.

### Immune system increases the volume of mitochondria in response to infection

We note that the body adjusts the number of white blood cells in circulation in response to infection. Chiu et al. showed that platelets produced in the bone marrow are supplied with more mitochondria, averaging 7.26 mitochondria per platelet when suffering from cancer verses 4.85 when healthy^70^. This implies that the body is actively increasing or decreasing the number of mitochondria in the immune system in response to infection. More numerous mitochondria in platelets yield more mitlets in circulation, which then transfer more mitochondria to the white blood cells to fight off infection. Thus, the body is systematically supplementing the number of mitochondria in white blood cells to match the demands of a given infection. Presumably, although this has not been described, one or more hormones or cytokines up-regulates the mitochondrial quantity per platelet.

### Mitochondria in white blood cells “burned out” by the stress of infection

According to Picard^71^, stressful conditions cause increased energy demands and mitochondrial DNA degeneration. Mitochondria serve many different functions, not only energy production but intracellular communication, inflammation, triggering apoptosis, and much more.^71^ Our findings show that immune cells quickly and preferentially absorb mitlets for extra mitochondria needed to support the system’s function, especially when fighting infection. This theory might be seen as analogous to a system of supply trucks providing soldiers in battle with more ammunition, supplies, and fuel, increasing these deliveries as needed to match the severity of the battle. Platelets serve many functions in the immune system; now, we show that they also deliver energy supplies to other cells.

### Mitlets yield measurable changes, whereas mitochondria isolated from the liver do not

Results confirm that the mitlets are much more effective than mitochondria isolated from the liver. We propose that mitlets are pre-designed and engineered by evolution to perform these duties. They are encased in extracellular vesicles containing various receptor patterns, causing them to be preferentially targeted to and quickly accepted by the white blood cells floating nearby in the blood.^11^ Mitochondria isolated from livers have no such vesicle and receptor coating, are not quickly absorbed by nearby immune cells, randomly flowing through the bloodstream to eventually become absorbed by other tissues such as lung, liver, or brain.^77,78^

### Many types of mitlets in various tissues

Other studies have shown that the immune system, in fact, has many different varieties of mitlets with different receptor patterns on the surface.^73^ Therefore, we might say that the immune system has a tool kit of different mitochondrial-bearing mitlets. These mitlets have also been shown to decline with age.^73, 76^ The brain also has neural-specific mitlets that transfer mitochondria between astrocytes and neurons^16,17^ and a similar system to serve cardiac cells.^75^ We note that cardiac cells, neurons, and white blood cells are all high-priority, mission-critical cells for the body. In case of fight-or-flight, escape from predators, or infections from constant injuries, the energy production efficiency of the mitochondria in these cells are critical to the organism’s survival. It makes logical sense that these cells have evolved to offload lower-priority mitochondrial production and maintenance activities to other support cells nearby, in order to conserve all available energies for survival. This would provide a significant competitive advantage. Thus, we may logically identify mitlets as a pervasive, evolutionarily-conserved subsystem of the body, which distribute scarce energy resources between cells and tissues in response to environmental conditions. Similar mitochondrial transfer occurs via stem cells and directly from cell-to-cell within tissues.^21^ Researchers are only now beginning to uncover this extraordinarily sophisticated web of energetic distribution, which is the fundamental basis of complex life.^79^

### The benefit of transplantation is temporary

From these results we see the mitlet transplant’s survival benefits decline steadily after the initial three-day infusion. This is consistent with our understanding of white blood cells. First, many white blood cells have short life spans. Thus, the beneficial effect of transplanted mitochondria will slowly wear off over a few days as the white blood cells undergo apoptosis and are replaced by wild-type white blood cells lacking the transplanted mitochondria. Additionally, according to Picard’s theory of mitochondrial damage due to stressful conditions^71^, we would expect that the freshly-transplanted younger mitochondria might quickly age and decay as their DNA is damaged by stressful usage. In essence, the young mitochondria quickly become old, in the heat of battle. We aim to explore this process in future experiments by performing additional injections on days 5, 7, 9 to determine if a continuous infusion of fresh replacement mitlets will allow the animals to survive longer.

### Potential for permanent regeneration of the immune system

It is interesting to note that the concept of exhaustion in immunotherapy is already a well-known issue in other types of immunotherapies, such as CAR-T or CAR-NK therapy.^82^ Transplantation of mitochondria via mitlets may also potentially apply to other types of immune senescence in the thymus or bone marrow, which also absorb mitlets, although in less proportion to white blood cells.^32^ For that reason, we hypothesize that with changes to the administration or dosing of mitlets, it might be possible to transplant large numbers of young mitochondria into solid immune tissues so that the young mitochondria become templates for mitochondrial replication and distribution to future generations of platelets and white blood cells. Thus, age-reversal of the immune system might potentially be permanent instead of temporary. This would contribute further to the health of the aged individual, potentially reducing inflammation or reducing the occurrence of cancer and other immune-mediated diseases of aging.

#### Limitations of the study

Although our Kentucky sepsis experiments showed significant improvements in survival and reductions in bacterial count, results from the Bristol H1N1 influenza experiments did not show statistical significance, although there was sufficient improvement to infer the effect, especially since it matched closely to the results from the sepsis experiments. The lack of significance in the H1N1 results is likely due to a low dosage of administration, the low number of animals, and, most importantly, the fact that donor and recipient animals were the same age. As we noted earlier, same-age donations were not expected to show strong results. A replicate study may be conducted for the Bristol experiment, using a larger number of animals and higher doses, to achieve statistical significance. The Kentucky sepsis experiments did not include positive controls, such as an anti-bacterial/anti-inflammatory treatment which is often used in sepsis tests^66^, nor did they include tests with same-age mice to provide context for the survival results, nor did they include dosing studies. These omissions were deemed acceptable in the short term since the tests are not intended to address sepsis treatments specifically but use Sepsis only as a convenient biomarker of immune system age. Nonetheless, these issues are planned for future experiments.

#### Prospects for future experimentation

Future tests should address safety, dosing, and improved controls. Tests in larger animals or humans will encounter many potential challenges, including increases in inflammatory mediators, conflicts between mitochondrial haplotypes or vesicle antigens, organ-specific challenges, effects from storage of mitochondria, auto-immune responses, high concentrations of calcium ions, routes of administration, and overall safety concerns. Assuming these challenges can be addressed, there seems no fundamental reason that these treatments could not be used in humans, for example, elderly patients with sepsis, influenza, COVID-19, some types of cancers, other infectious diseases, and soldiers with battlefield injuries or sepsis. Another potential target population would be children with mitochondrial mutation diseases such as Leigh Syndrome, who have weakened immune systems and can suffer very negative outcomes from common infections. These children might be given mitlet transplants to temporarily make their immune systems “normal” to fight infection.^63,64,65^

Blood banks, which already have platelet transfusions regularly, might isolate mitlets, refrigerate them, and use them as needed in exceptional cases. However, it seems unlikely that the number of platelets currently available would create enough mitlets to treat many patients. Just as donor livers or kidneys are in short supply, we might expect donor mitochondria to be chronically short as well. Mitochondrial transplantation requires large volumes; small numbers confer little advantage. Therefore, developing a bioreactor manufacturing process that could grow mitlets (mitochondria encased in extracellular vesicles or an equivalent coating, along with artificial receptors to target specific tissue types) would be more practical for mass-market distribution and global use. Mitrix Bio, the sponsor of this study, has an active program to build this mitochondrial bioreactor.^74^

#### Potential value to society

If this process is found to be safe and efficacious in larger animals and humans, then it would represent a new type of immunotherapy with broad application to human medicine and pandemic response. We would term this HISET therapy (Human Immune System Energetics Transplant). HISET therapy could potentially serve as a “general purpose” immune system booster, which could be administered in combination with other antiviral or anti-bacterial treatments. Many elderly patients with infectious diseases end up in the hospital or the Intensive Care Unit (ICU), both of which are high-cost and a significantly drain on national healthcare budgets. By boosting the immune system of elderly patients with mass-manufactured transplantation-quality mitochondria, even if such a treatment is temporary, we could help large numbers of patients avoid dangerous and expensive ICU stays. These treatments might also apply to chronic diseases and pandemic management. For example, studies have shown that COVID-19 can cause significant mitochondrial damage and might contribute to Long Covid.^80,81^ Therefore, mitochondrial transplantation therapy has potential for chronic post-infection diseases such as Long Covid, Long Sepsis, or post-chemotherapy.

## CONTRIBUTIONS

HS, SP, AA, and SS designed and conducted the experiments. TB wrote the manuscript. BA and VM edited and consulted on the manuscript.

## ETHICS STATEMENT

All experimental procedures at the University of Kentucky were approved by the Institutional Animal Care and Use Committee. Animal handling techniques in these studies were performed as described and approved in our Animal Use Protocol #2009-0541 and were in accordance with the National Institutes of Health guidelines for ethical treatment. All experimental procedures at Charles River Labs in Bristol, England, were approved by the Institutional Animal Care and Use Committee.

---

**FIGURE 1.**
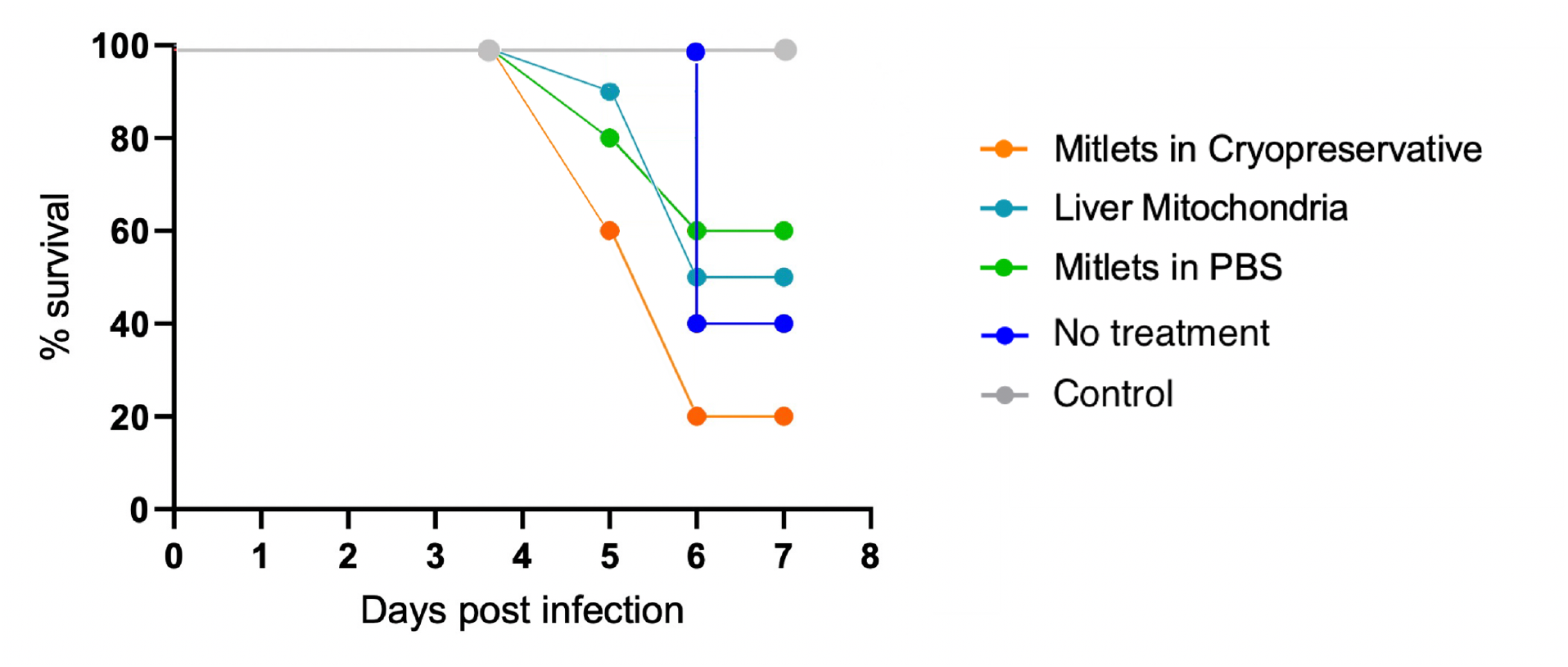
Platelet extracellular vesicles (mitlets) isolated from young mice and transplanted into same-age recipient mice with H1N1 influenza infection, improved survival. Kaplan-Meier Survival Graphs. Some data points have been nudged for clarity. “Naïve control” received no H1N1 dosing. “Vehicle control” received the H1N1 dosing but was given no treatment. Others received the H1N1 dosing and were given mitlet or mitochondria as shown in legend. Improvement did not reach statistical significance.

**FIGURE 2.**
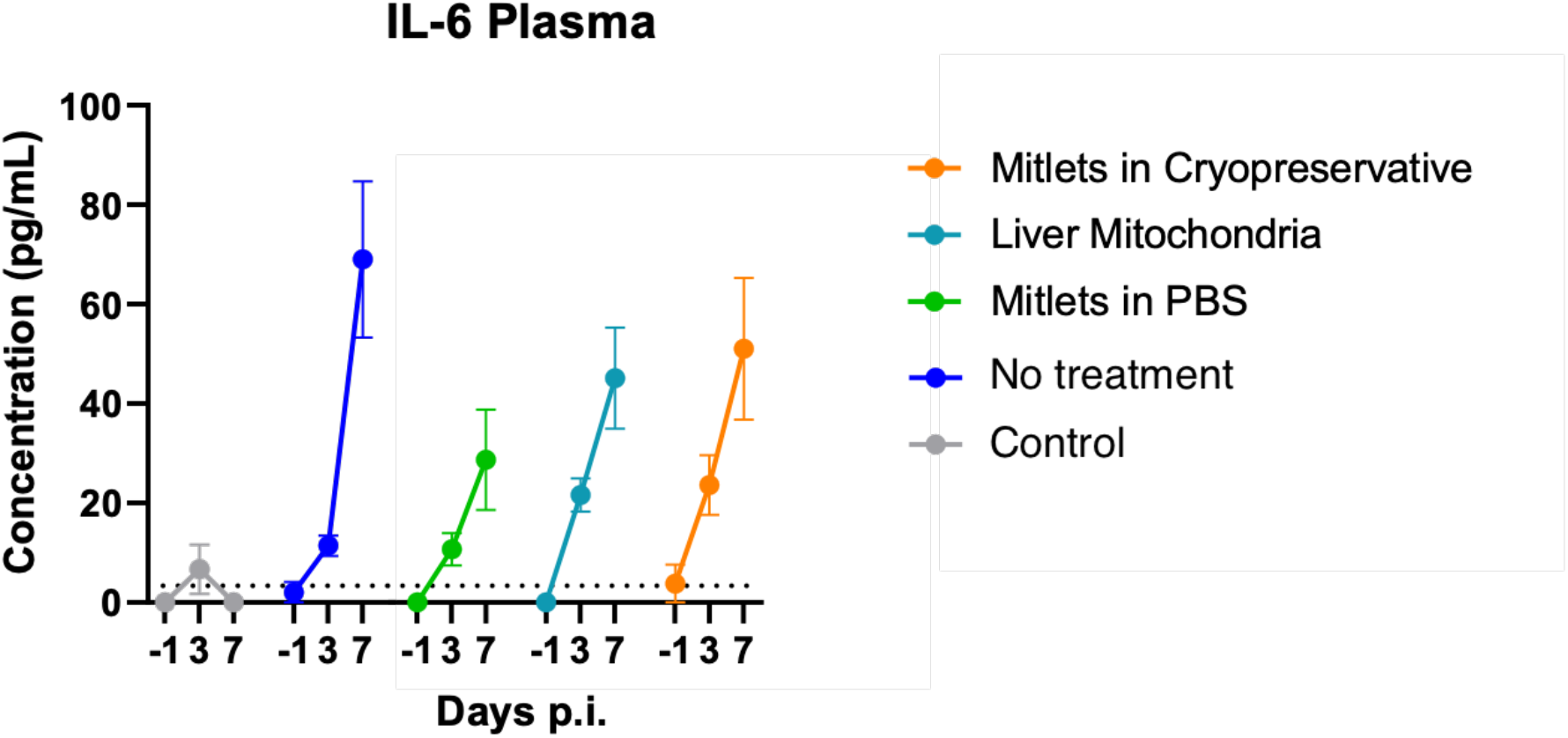
Platelet extracellular vesicles (mitlets) isolated from young mice and transplanted into same-age recipient mice with H1N1 influenza infection, reduced cytokine levels (plasma IL-6), day -1 to day 7. Blood was collected from each animal and processed to plasma for cytokine analysis by Luminex (n=5-10 per group). Samples were analyzed in singlicate at a 1 in 4 dilution. Each dot represents the group mean + SEM at each time point. The functional Lower Limit of Quantification (LLOQ) (3.34 pg/mL) is plotted as a dotted black line. Values <LLOQ are plotted as 0 pg/mL. Statistical significance analysis was attempted using a Mixed-effects model, however due to values <LLOQ a p value could not be calculated. “Naïve control” received no H1N1 dosing. “Vehicle control” received the H1N1 dosing but was given no treatment. Others received the H1N1 dosing and were given mitlet or mitochondria as shown in legend. No results reached statistical significance.

**FIGURE 3.**
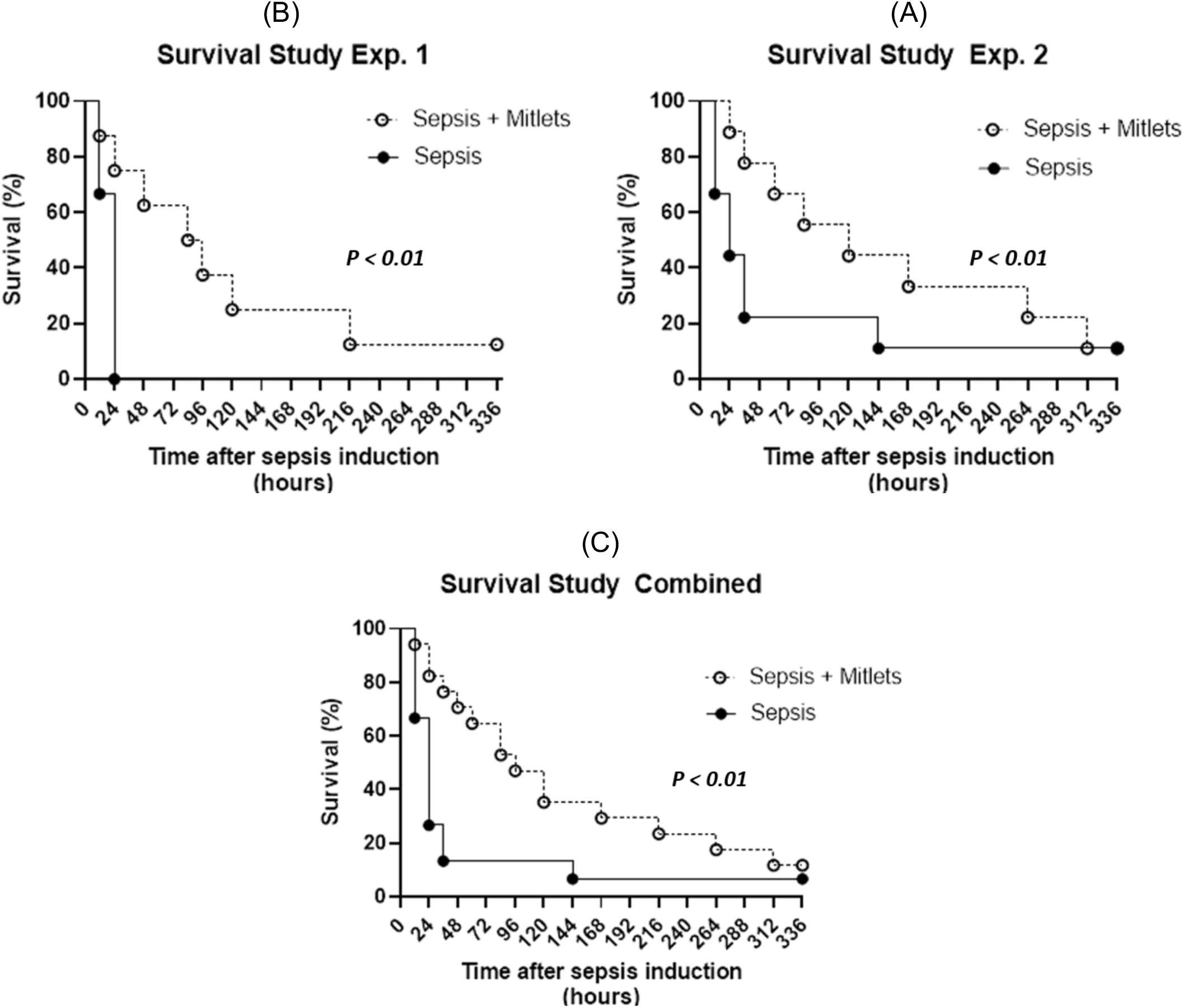
Platelet extracellular vesicles (mitlets) isolated from young mice and transplanted into aged recipient mice with sepsis infection significantly improved survival. (A) Mitlet injections, first experiment. (B) Mitlet injections, second experiment. (C) Mitlet injections, first and second experiments combined. Statistical significance as shown.

**FIGURE 4.**
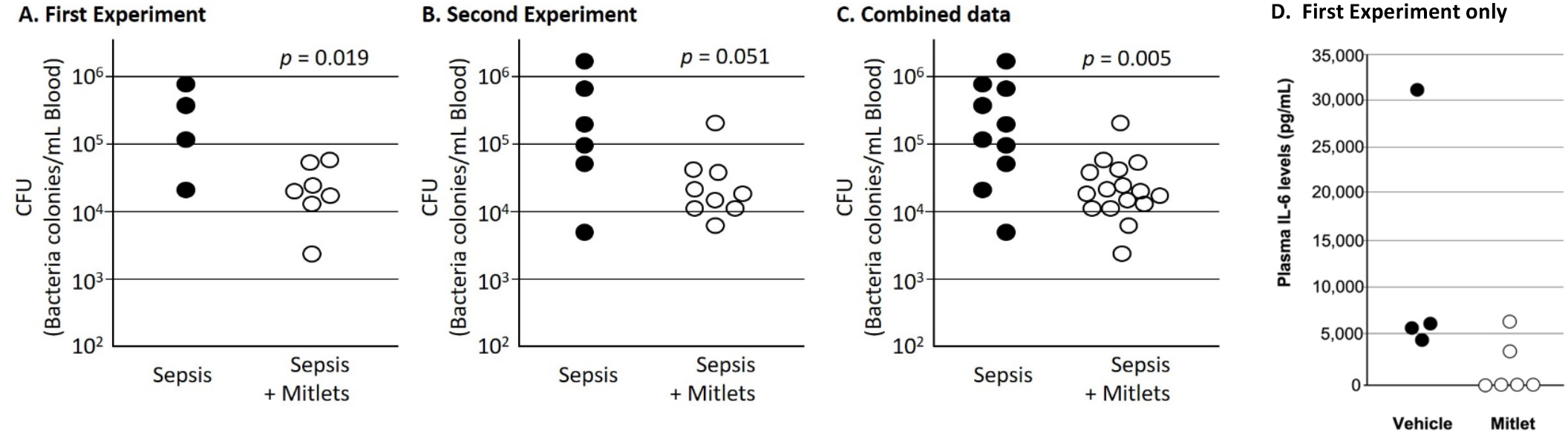
(A/B/C/D) Platelet extracellular vesicles (mitlets) isolated from young mice and transplanted into aged recipient mice with sepsis infection, significantly reduced cytokine IL-6 and bacterial levels. (A) Bacterial levels after mitlet injections, first experiment. (B) Bacterial levels after mitlet injections, second experiment. (C) Bacterial levels after mitlet injections, first and second experiments combined. (D) Plasma IL-6 levels for first experiment only. Liver isolated mitochondria not shown. Statistical significance as shown.

**FIGURE 5.**
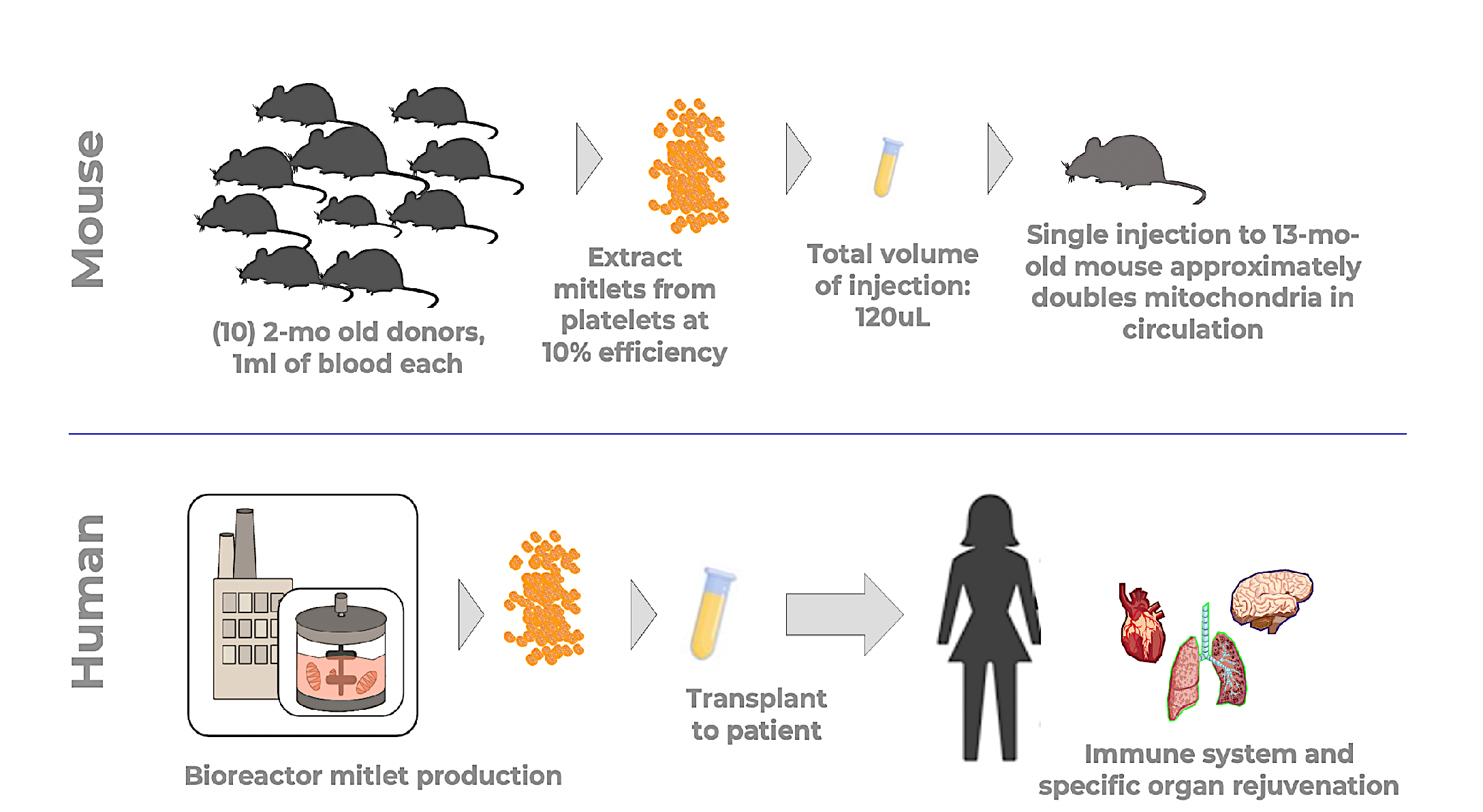
Schematic. (Mouse) Approximately 10mL of donor mouse blood process into mitlets, yielding 120uL of injectate per 1 injection. (Human) Proposed for future – bioreactor facility produces large volumes of allogenic or autologous mitochondria coated with protein coating and receptor molecules, simulating function of naturally-occurring mitlets.

